# Cardiomyocyte hypertrophy co-localizes with accumulation of hyaluronan in cMyBPC^+/-^ mice hearts

**DOI:** 10.64898/2026.01.30.702716

**Authors:** Anna Strnadová, Madelene Ericsson, Rakel Nyrén, Ilona Dudka, Stellan Mörner, Urban Hellman

## Abstract

Hypertrophic cardiomyopathy (HCM) is a genetic heart disease characterized by left ventricular hypertrophy. Mutations in genes for sarcomere proteins, e.g. in cardiac myosin-binding protein C (cMyBPC), cause symptoms like heart failure and sudden cardiac death. In HCM, the heart utilizes high amounts of glucose. Increased glucose metabolism creates intermediates for hyaluronan (HA) synthesis, resulting in HA accumulation within the extracellular space, which could contribute to cardiomyocyte hypertrophy.

We aimed to describe the connection between HA and cardiomyocyte hypertrophy using cMyBPC^+/-^ mice treated with two different β-blockers – either metoprolol or propranolol – for 11 months.

The results showed that HA metabolism is altered in HCM, and consequently, HA is significantly more abundant in hearts of cMyBPC^+/-^ mice compared to wild-type mice. High HA abundance correlated with increased cardiomyocyte size. Gene expression of HA synthase 2 was elevated in cMyBPC^+/-^ mice. Treatment with propranolol significantly increased HA synthase 3 expression, while HA synthase 2 was maintained at wild-type level. Treatment with metoprolol or propranolol did not prevent HA accumulation nor decreased cardiomyocyte hypertrophy. These findings show a novel connection between high HA abundance and cardiomyocyte hypertrophy in cMyBPC^+/-^ mice.

## Introduction

Hypertrophic cardiomyopathy (HCM) is a genetic disease affecting the cardiac muscle, often caused by heterozygous mutations in genes encoding sarcomere proteins such as myosin heavy chain or cardiac myosin binding protein C (cMyBPC). Affecting primarily the left ventricle, this condition causes hypertrophy, predominantly of the interventricular septum. This may lead to hypercontractility, left ventricular outflow tract obstruction, diastolic dysfunction, and arrhythmias. The prevalence of HCM in the general population is 0.2%, with males being more commonly affected than females, although the inheritance pattern is autosomal dominant [1–3]. Individuals with this condition can begin to experience symptoms at any point in their lives, from newborns to older adults. HCM is a common cause of sudden cardiac death, which may occur as the first manifestation of the disease [4,5]. Treatment consists of either pharmaceutics designed to relieve symptoms of heart failure and arrhythmias, such as β-blockers, or invasive surgical procedures [6,7]. There are no therapies available to prevent the onset of HCM in humans [2,7]. However, in children with HCM, hypertrophy can be attenuated by high doses of β-blockers, such as propranolol [8,9].

Recent research suggests that myocardial energetic pathways are impaired in HCM; sarcomere gene mutations lead to inefficient ATP utilization and mitochondrial dysfunction [10]. It has previously been shown that in HCM the expression of genes connected to energetics is decreased [11]. This results in unmet high energy demands in the heart, causing energetic stress and adverse remodelling [11,12]. To keep up with the elevated energy demands, the heart begins to use glucose as its primary source for ATP production as it requires less oxygen. In contrast, healthy hearts generate ATP mostly through β-oxidation [12,13]. The involvement of glucose metabolism in HCM pathogenesis is not yet properly understood. Nonetheless, an example of high blood glucose-caused cardiac hypertrophy can be seen in infants born to diabetic mothers, where the children suffer from severe cardiac hypertrophy at birth; however, this condition resolves spontaneously as normoglycemia establishes during the following months [14].

On a microscopic level, HCM hearts contain hypertrophied cardiomyocytes, cellular disarray and fibrosis. A previous study [15] reported that hearts from HCM patients show high variability – not only between individuals, but also within sections from the same heart. They have demonstrated that severely re-modelled and hypertrophied cardiomyocytes neighbour normally sized and aligned cardiomyocytes, creating focal patches of pathological and healthy tissue throughout the heart.

In a previous study from our lab, we described a novel link between elevated glucose usage and the development of cardiac hypertrophy; our results also suggested an increased risk of arrhythmia and sudden cardiac death in HCM. The connection between glucose metabolism and cardiac hypertrophy was observed both in cardiac tissue from human HCM patients, and in cultured rat cardiomyocytes, as well as in rats with induced cardiac hypertrophy [16–19]. We observed that increased glucose metabolism leads to increased synthesis of the glycosaminoglycan hyaluronan (HA), which is present in the extracellular matrix of the heart [16,17,19,20]. Excessive deposition of HA in the heart might therefore contribute to arrhythmias and fibrosis, as well as increased cardiomyocyte size. HA is a non-sulphated glycosaminoglycan consisting of repeating disaccharide units of glucuronic acid (GlcA) and N-acetyl-glucosamine (GlcNAc). It is one of the most abundant components of the extracellular matrix, where it plays a crucial role in tissue hydration, cell survival and cell migration [21].

In the focal patches of abnormal cardiac tissue, the increased glycolysis leads to an accumulation of glycolytic intermediates, which are further utilized for synthesis of HA. An excessive amount of this glycosaminoglycan in the extracellular space could affect cell-to-cell communication as well as electrical conduction through the heart. HA accumulation could interfere with proper heart function, contributing to hypertrophy and a further increase of glycolysis, creating a vicious pathogenic circle. Furthermore, overabundant HA, mainly fragmented low molecular weight molecules, around individual cardiomyocytes can contribute to inflammation [22]. A way to affect glucose metabolism, and possibly by extension HA metabolism and hypertrophy, is by administering β-blockers; this commonly used therapy for cardiac dysfunction is known to decrease cellular glucose uptake, insulin sensitivity and hepatic glucose production [23,24].

Here, we focused on exploring the connection between HA, glucose and hypertrophy using cMyBPC^-/+^ mice, which are described to develop left ventricular hypertrophy akin to human HCM at around 11 months of age [25]. To investigate the effect of these two treatments on HA and cardiac hypertrophy, the mice were treated from weaning with either high-dose propranolol, commonly used for treating children with HCM [8,26], or the standard dose of metoprolol, predominantly used in the clinics to treat adults with HCM [2,27].

## Materials and methods

### Animal model

Male heterozygotic for cardiac myosin binding protein-C mice (cMyBPC^+/-^) [25] and male wildtype (WT) C57BI/6J mice were included in the study. The animals were generated in-house by breeding heterozygous males with WT female mice, to ensure healthy gestation and maternity. The pups were genotyped and assigned to groups according to their genotype. The first study with metoprolol was performed three years prior to the propranolol study. An overview of included mice and treatments is summarized in Fig 1.

**Fig. 1.**
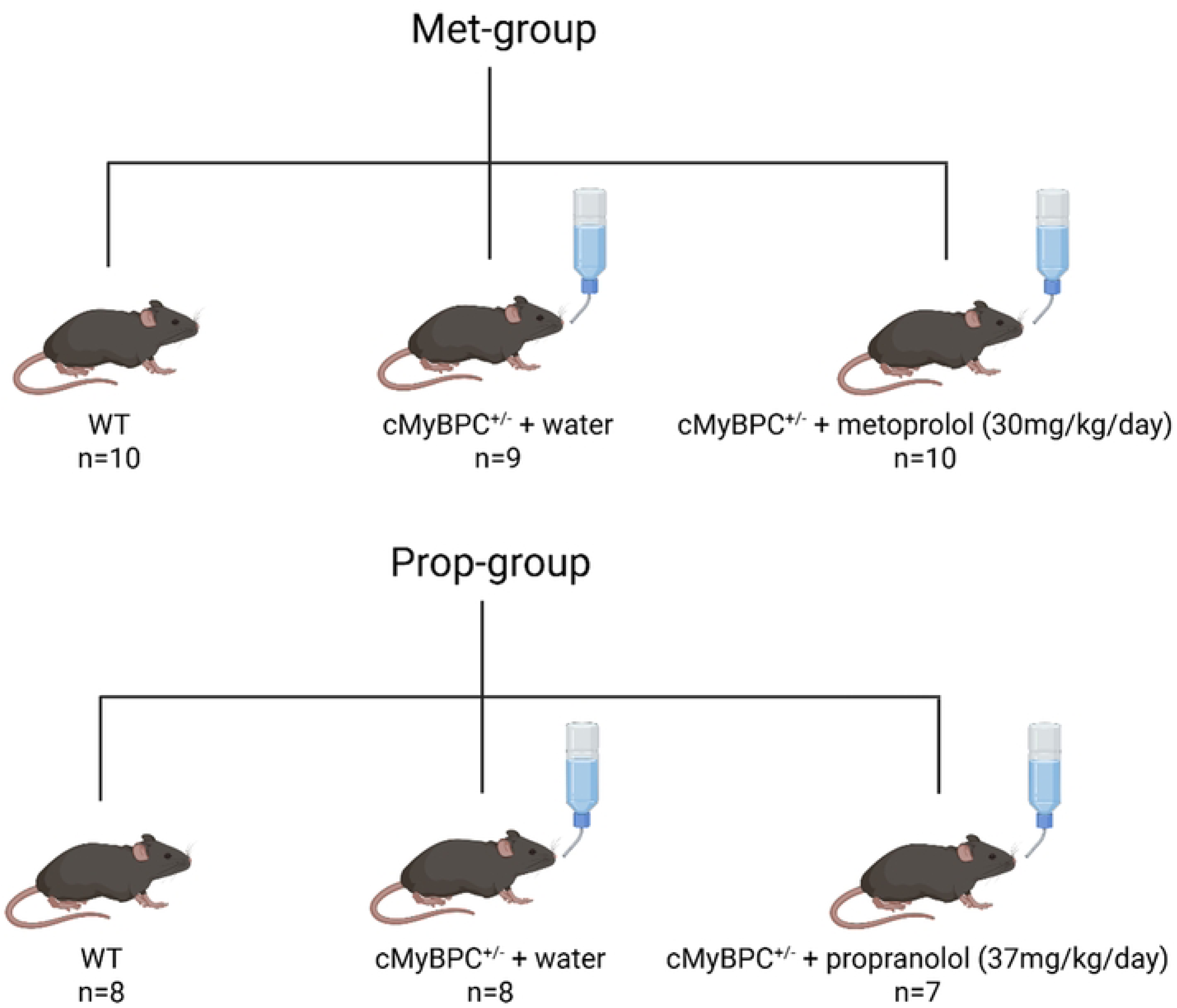
Overview of mice included in this study. Male wildtype (WT) C57BI/6J mice and male heterozygous cardiac myosin binding protein-C (cMyBPC^+/-^), given either pure drinking water, or β-blocker dissolved in their drinking water; metoprolol (30mg/kg/day) in Met-group (cMyBPC^+/-^+ met mice) or propranolol (37mg/kg/day) in Prop-group (cMyBPC^+/-^+ prop mice).

The mice were housed in groups of 2-4 mice/cage under standard housing conditions (12:12h light-dark cycle, 22°C temperature and 45-50% humidity), with free access to standard rodent chow and drinking water. Treatment started after weaning. cMyBPC^+/-^ mice were given either pure drinking water or a β-blocker dissolved in their drinking water (cMyBPC^+/-^+met mice given metoprolol (30 mg/kg/day) in Met-group or cMyBPC^+/-^+prop mice given propranolol (37 mg/kg/day) in Prop-group) (Fig 1). Blood glucose was measured at endpoint from tail vein blood using a hand-held ACCU-Chek AVIVA device (Roche Diagnostics, Solna, Sweden). When twelve months old, the mice were sacrificed via cervical dislocation. Hearts were excised immediately after euthanasia, and dissected into specific regions (left ventricle, right ventricle, apex and atria) snap-frozen in liquid nitrogen and stored at -80°C until use.

The mice were cared for and used in accordance with Swedish Animal Welfare Act and approved by the Animal Review Board at the Court of Appeal for Northern Norrland in Umeå, Sweden (A62/14).

### Echocardiography

Cardiac function was assessed using the Vevo 2100 (VisualSonics, Fujifilm, Toronto, Canada), and the MS500D transducer. The examination was performed during light isoflurane anaesthesia (1.0–2.0% in 800 ml/min O_2_) (Baxter Medical, Kista, Sweden). The level of anaesthesia was adjusted to keep the respiration rate at 90–110 breaths/min [46]. Image acquisition was previously described [46,47]. Image analysis was performed off-line in a blinded manner by AS, using the Vevo LAB 5.10.0 software (VisualSonics, Fujifilm). The mean of three strokes was used for each parameter and mouse.

### Positron Emission Tomography (PET) for glucose uptake

To evaluate cardiac glucose uptake in the Prop-group, the radioactive glucose analog [18F]-Fluorodeoxyglucose (FDG) was administered. In short, during light isoflurane anesthesia, as described under echocardiography, mice were injected with 70-100µL (12.8±4.8MBq) of clinical grade FDG in saline, prepared at the Nuclear Department at Norrlands University Hospital, Umeå, Sweden. A tailor made 27G needle and catheter was used to ensure correct administration in the tail vein. After injection, mice were allowed to be awake and freely moving in their cage. After 120 minutes, mice were again sedated with isoflurane and CT was performed for gross morphological orientation and then scanned for a 10-minute static PET acquisition (nanoScan PET/CT, Mediso, Hungary). Images were reconstructed over a 98 mm axial field-of-view to a 0.4 x 0.4mm slice resolution, employing a 3D iterative reconstruction, four iterations and four subsets (Mediso Tera-Tomo 3D). The reconstruction was performed with delayed-window random correction, spike filter, scatter, and CT-based attenuation corrections. Tracer uptake in myocardial tissue was evaluated by using imlook4d (https://github.com/JanAxelsson/imlook4d), and quantified as standardized uptake values (SUV), using the formula: *SUV = C / (i / m)*; with *C* being the measured tissue activity concentration (Bq/mL) in the image, *i* the injected dose, and *m* the mouse body weight. After PET/CT scan, mice were allowed to wake up and return to their home-cages. FDG-PET was only performed for Prop-group, as it was not available when Met-treatment was performed.

### RT-qPCR

Collected tissues were snap frozen in liquid nitrogen and stored at -80°C. The apex of the heart was used for gene expression analyses. Tissues were thawed in RNA*later*^TM^-ICE (Invitrogen, ThermoFischer; AM7030, Waltham, USA) according to manufacturer’s protocol. Total RNA was extracted using RNeasy Mini Fibrous Kit (Qiagen; 74704; Hilden, Germany). All samples were treated with RNase-Free DNase (Qiagen; 79254). RNA concentration and purity were measured on a NanoDrop. cDNA was prepared with RevertAid H minus Reverse Transcriptase (Thermo Scientific) and Random Hexamer primer (Thermo Scientific) in a Thermocycler from Biometra.

MasterMix TaqMan Fast Advanced (Applied Biosystems, ThermoFischer; 4444557), housekeeping gene (GAPDH-VIC or Rpl32-VIC) and one of the following genes (Table 1) were pipetted onto the MicroAmp Optical 96-Well plates (Applied Biosystems, ThermoFischer; 4306737) and cDNA of each sample was added to a final concentration of 40 ng/well. All samples were run in triplicates. For negative control, cDNA was replaced by nuclease-free water (Fisher Bioreagens; BP2484-50). ROX was used as a passive reference.

**Table 1.** Gene probes used for RT-qPCR.

| **Assay/gene** | **Protein name** | **Cat. No.** |
| --- | --- | --- |
| ***Has1*** | Hyaluronan synthase 1 | Mm03048195_m1 |
| ***Has2*** | Hyaluronan synthase 2 | Mm00515089_m1 |
| ***Has3*** | Hyaluronan synthase 3 | Mm00515092_m1 |
| ***Hyal1*** | Hyaluronidase 1 | Mm00476206_m1 |
| ***Hyal2*** | Hyaluronidase 2 | Mm01230688_g1 |
| ***Cemip*** | Cell migration inducing hyaluronidase 1 | Mm00472921_m1 |
| ***Tmem2*** | Transmembrane protein 2 | Mm00459599_m1 |
| ***Slc2a4/Glut4*** | Glucose transporter 4 | Mm00436615_m1 |
| ***Gapdh*** | Glyceraldehyde-3-phosphate dehydrogenase | Mm99999915_g1 |
| ***Rpl32*** | 60S ribosomal protein L32 | Mm02528467_g1 |

Gene expression was quantified using QuantStudio 6 Flex System instrument. The amplification was run in comparative mode and analysed in QuantStudio 6 software (v1.7.2). Relative quantification was calculated according to the 2^-ΔΔCT^ method using either GAPDH or Rpl32 as a housekeeping gene and WT mouse sample as the control group.

### Histopathological evaluation and scoring of cardiac fibrosis

Histopathological assessments were performed on 4 µM formalin fixed and paraffin embedded sections from the left ventricle of each mouse [48]. Screening for histopathological changes such as disarray was done using standard hematoxylin and eosin staining. Collagen fibres were visualized using Masson’s trichrome stain (HT15, Sigma-Aldrich, St Louis, MO, USA) according to the manufacturer’s protocol. The presence of interstitial fibrosis was evaluated in a blinded manner by RN in twenty randomly selected high power fields (0.24 mm^2^) per heart using a Nikon Eclipse Ci light microscope. Each area was assessed using a semi-quantitative scoring system (0 = no fibrosis, 1 = slight, 2 = moderate, 3 = severe) and presented as mean value per individual [49].

### Hyaluronan quantification and cell size analysis

For hyaluronan staining, cardiac sections were pretreated with 3% H_2_O_2_ in methanol and 1% bovine serum albumin solution (10mg/ml in PBS) before overnight incubation at 4°C with biotinylated hyaluronic acid binding protein probe (100ug/ml in PBS, 1:40) (HABP, Sigma-Aldrich, St. Louis, USA).

The HABP probe was visualised using VECTASTAIN® Elite® ABC-HRP Kit (Vector Laboratories, Newark, USA) and DAB substrate kit (Vector Laboratories) according to manufacturer’s instructions. Cell nuclei were stained using Mayers HTX PLUS (Histolab Products AB, Askim, Sweden). The samples were imaged using Pannoramic 250 Flash III digital scanner (3DHistech, Budapest, Hungary).

Using QuPath software (version 0.5.1), each sample was annotated as a region of interest (ROI) using the pixel qualification function. Subsequently, every sample was divided into square tiles (200 µM a side) that were trimmed to ROI borders. This tile approach was implemented to mitigate the individual differences among samples.

To visualise hyaluronan staining, pixel qualification was performed (settings: resolution 0.24 µM/px, DAB channel, sigma 0.5, threshold 0.1). Next, in each sample, we identified tiles that contained no tissue edges, no excessive tearing and no vessels. Tiles containing vessels were excluded as vessels naturally contain high amounts of hyaluronan. This could distort the results as there is variation between the samples in how many vessels they contain. Furthermore, tiles with tearing or edges were also disqualified, as they contain a lot of empty space which would be incorrectly quantified as ‘no staining’ due to the software setup, thereby introducing errors into the analysis.

From tiles that fulfilled all criteria, 16 tiles containing high amounts of hyaluronan staining (high c[HA] areas; HA staining > 5%) and 16 tiled that contained little-to-no hyaluronan staining (low c[HA] areas; HA staining < 5%) were selected. In each of these tiles, we quantified the percentage of positively stained tissue, which corresponds to the amount of hyaluronan. For each sample, we could obtain an average percentage of positive staining in high c[HA] areas and in low c[HA] areas.

For cell width quantification, we again used QuPath software. Using the same tiles as for HA quantification, the width of 50 cells per sample was measured, approximately 3 per selected tile. The cell width was measured manually using the QuPath’s ‘Line’ function. For the cell to be considered appropriate for measurement, it had to meet the following selection criteria: 1) the cell had to be longitudinally cut, 2) the cell nucleus had to be clearly visible and symmetric, 3) the cell’s edges had to be clearly defined.

### Statistical analysis

Data analyses pertaining to comparing different experimental groups, such as one-way ANOVA or t-tests, were performed using GraphPad Prism statistical analysis package (version 10.4.1 (627)).

Pearson correlation analyses were performed using SPSS statistical package (version 29.0.2.0, BMI).

Data are expressed as mean ± SD. Statistical significance was set at P<0.05.

## Results

### Cardiac function and glucose uptake by FDG-PET

To evaluate cardiac function and pathological cardiac growth in cMyBPC^+/-^ mice, as well as possible effects of two different kinds of long-term β-blockade treatment, high-frequency echocardiography was performed. In the first experiment, cMyBPC^+/-^ mice were treated with metoprolol from weaning to 12 months of age (Met-group). The same experiment was repeated three years later, but a non-selective β-blocker propranolol was used instead (Prop-group) (Fig 1). As HCM results in cardiac hypertrophy, left ventricular dimensions were assessed at 12 months. However, neither anterior nor posterior wall hypertrophy was found in cMyBPC^+/-^ mice in either of the two studies. (Table 2). Nevertheless, in Met-group, all cMyBPC^+/-^ mice had significantly reduced stroke volume and cardiac output. In contrast, cMyBPC^+/-^ mice in the Prop-group exhibited similar measurements as WT mice. Overall, none of the β-blockers appeared to influence any of the parameters measured by echocardiography.

**Table 2.** Echocardiographic data summary at 12 months of age.

| Echocardiography | Met-group |  |  | Prop-group |  |  |
| --- | --- | --- | --- | --- | --- | --- |
|  | WT | cMyBPC<sup>+/-</sup> | cMyBPC<sup>+/-</sup> + met | WT | cMyBPC<sup>+/-</sup> | cMyBPC<sup>+/-</sup> + prop |
|  | (n = 7) | (n = 9) | (n = 9) | (n = 5) | (n = 6) | (n = 4-6) |
| HR (bpm) | 482 ± 33.8 | 468 ± 25.0 | 454 ± 47.0 | 437 ± 83.8 | 459 ± 65.3 | 421 ± 45.0 |
| SV (μL) | 21.6 ± 1.6 | 16.5 ± 2.2*** | 17.4 ± 2.6** | 29.2 ± 5.4 | 31.9 ± 3.9 | 30.0 ± 4.7 |
| CO (mL/min) | 10.4 ± 1.1 | 7.6 ± 1.2*** | 7.8 ± 0.9*** | 12.5 ± 2.4 | 14.4 ± 1.7 | 12.5 ± 1.7 |
| ESV (μL) | 39.8 ± 4.4 | 38.4 ± 5.8 | 38.4 ± 6.9 | 40.8 ± 10.2 | 43.3 ± 4.9 | 41.2 ± 3.7 |
| EDV (μL) | 61.4 ± 4.2 | 54.0 ± 5.9 | 55.7 ± 8.1* | 70.0 ± 10.1 | 75.1 ± 4.4 | 71.2 ± 7.0 |
| FS M-mode (%) | 18.2 ± 1.8 | 16.9 ± 3.7 | 18.8 ± 5.1 | 22.1 ± 4.6 | 20.0 ± 2.7 | 22.4 ± 1.1 |
| EF M-mode (%) | 37.7 ± 3.2 | 35.2 ± 7.0 | 38.6 ± 9.0 | 44.4 ± 8.1 | 40.8 ± 4.8 | 45.1 ± 1.8 |
| AWd (mm) | 0.66 ± 0.05 | 0.70 ± 0.07 | 0.68 ± 0.09 | 0.58 ± 0.05 | 0.71 ± 0.05 | 0.60 ± 0.05 |
| AWs (mm) | 0.86 ± 0.03 | 0.87 ± 0.1 | 0.90 ± 0.12 | 0.61 ± 0.07 | 0.72 ± 0.07 | 0.57 ± 0.05 |
| LVIDd (mm) | 4.68 ± 0.09 | 4.48 ± 0.27 | 4.52 ± 0.26 | 4.75 ± 0.35 | 4.82 ± 0.15 | 4.63 ± 0.24 |
| LVIDs (mm) | 3.83 ± 0.11 | 3.75 ± 0.25 | 3.66 ± 0.40 | 3.71 ± 0.48 | 3.85 ± 0.20 | 3.60 ± 0.19 |
| PWd (mm) | 0.84 ± 0.03 | 0.80 ± 0.10 | 0.86 ± 0.10 | 0.77 ± 0.05 | 0.80 ± 0.05 | 0.80 ± 0.10 |
| PWs (mm) | 0.99 ± 0.05 | 0.92 ± 0.09 | 1.01 ± 0.14 | 0.95 ± 0.10 | 0.92 ± 0.10 | 1.00 ± 0.10 |

Data presented as mean ± SD. WT, wild type; cMyBPC<sup>+/-</sup> + met, cMyBPC<sup>+/-</sup> mice treated with metoprolol; cMyBPC<sup>+/-</sup> + prop, cMyBPC<sup>+/-</sup> mice treated with propranolol. FS, fractional shortening; SV, stroke volume; EF, ejection fraction; CO, cardiac output; ESV, end-systolic volume; EDV, end-diastolic volume; AW, left ventricular anterior wall dimension; LVID, left ventricular internal dimension; PW, left ventricular posterior wall dimension; d, diastole; s, systole. Measurements using the parasternal long axis in B-mode or M-mode. Analysed by one-way ANOVA with Tukey's post hoc test. Significance WT vs cMyBPC<sup>+/-</sup>: \*P<0.05, \*\*P<0.01, \*\*\*P<0.001. Significance WT vs cMyBPC<sup>+/-</sup> + met: \*P<0.05, \*\*P<0.01.

In both experiments heart weight/body weight ratio was similar between WT and cMyBPC^+/-^mice with no significant change after β-blocker treatment (Table 3). Blood glucose, measured at 12 months of age, did not differ regardless of genotype or treatment.

**Table 3.**
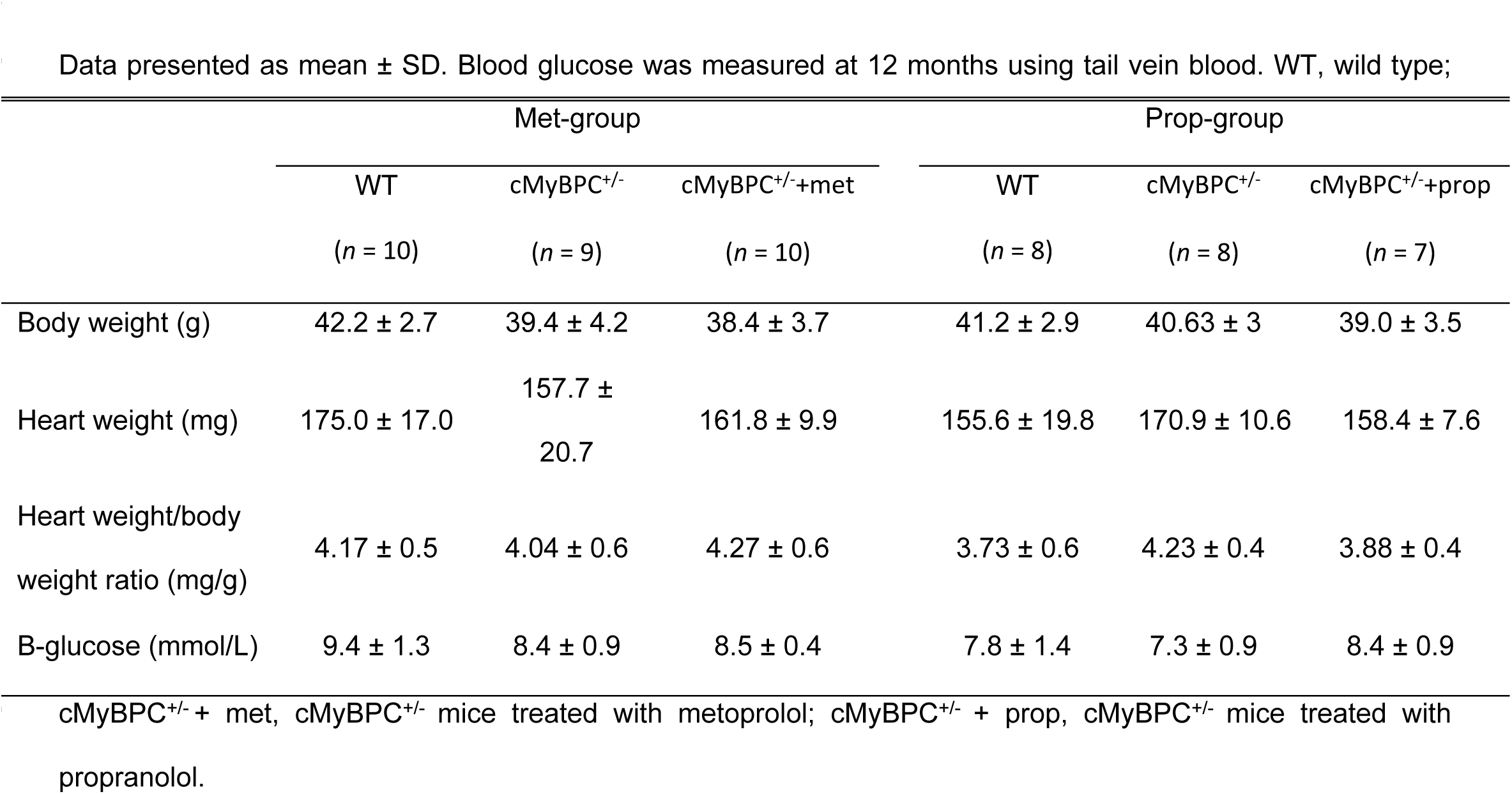
Gravimetrical data summary at 12 months of age.

To evaluate if cardiac tissue had an increased uptake of glucose *in vivo*, FDG-PET was used. Glucose uptake was only measured in Prop-group, as the technique was not available at the endpoint of Met-group. The cMyBPC^+/-^ and cMyBPC^+/-^+ prop mice showed a non-significant increase in glucose uptake compared to the WT mice. Propranolol treatment did not alter glucose uptake. Interestingly, cMyBPC^+/-^+ prop mice showed two groups of data points – one group with low uptake and one with high uptake, with no data at the mean (Fig 2).

**Fig 2.**
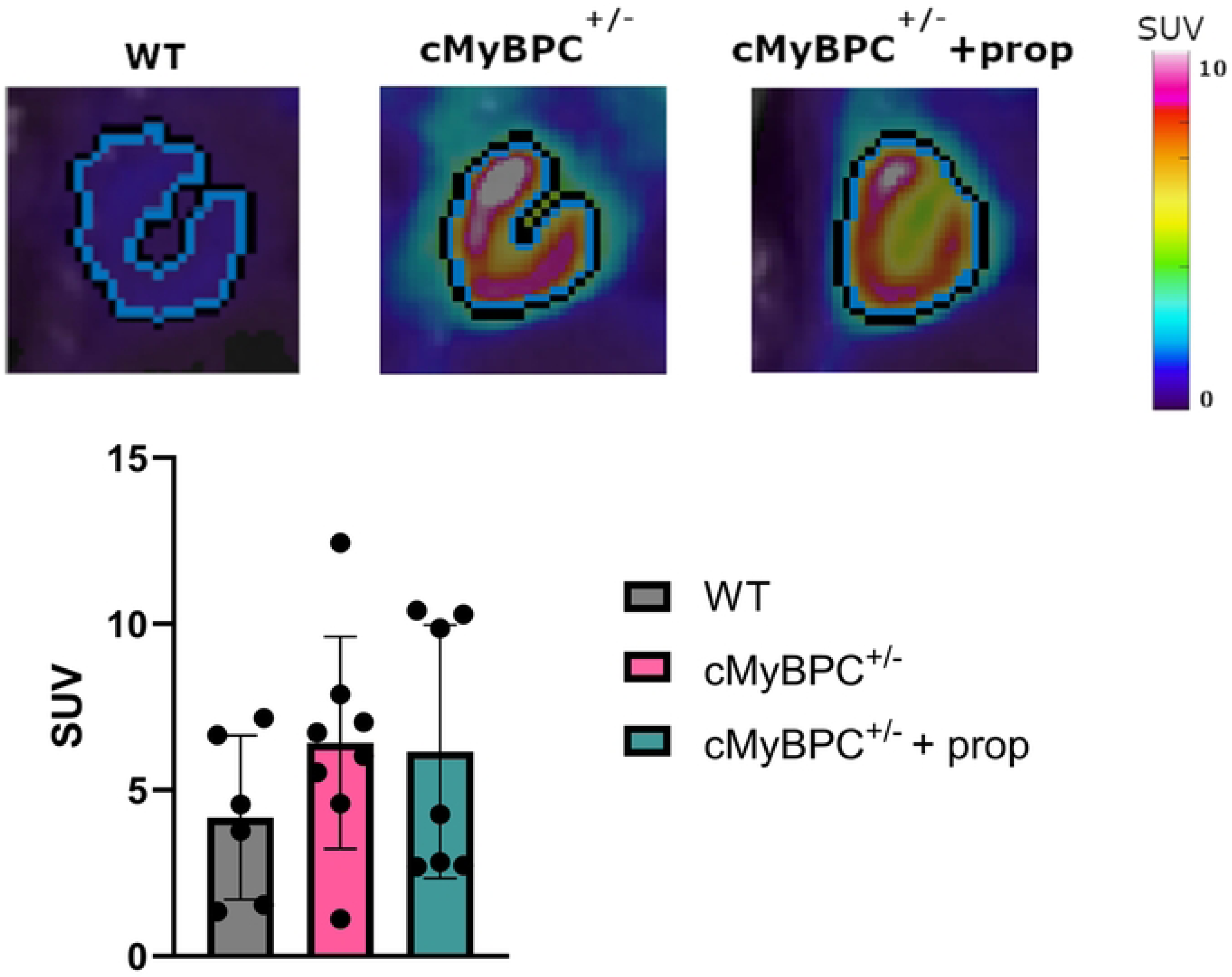
Maximal cardiac glucose uptake by FDG-PET presented as mean SUV ± SD. WT, wild type; cMyBPC^+/-^ + prop, cMyBPC^+/-^ mice treated with propranolol. SUV = standard uptake value.

### Histology

General histopathological assessment of cardiac sections stained with standard haematoxylin and eosin did not reveal any areas of disarray in the cMyBPC^+/-^ mice, one of the histopathological hallmarks of HCM. However, other signs of hypertrophy were found, such as enlarged, irregular and box-shaped cardiomyocyte nuclei. In addition, the cardiomyocytes in some areas of cMyBPC^+/-^ mice hearts appeared visually larger (Fig 3, upper panel).

**Fig 3.**
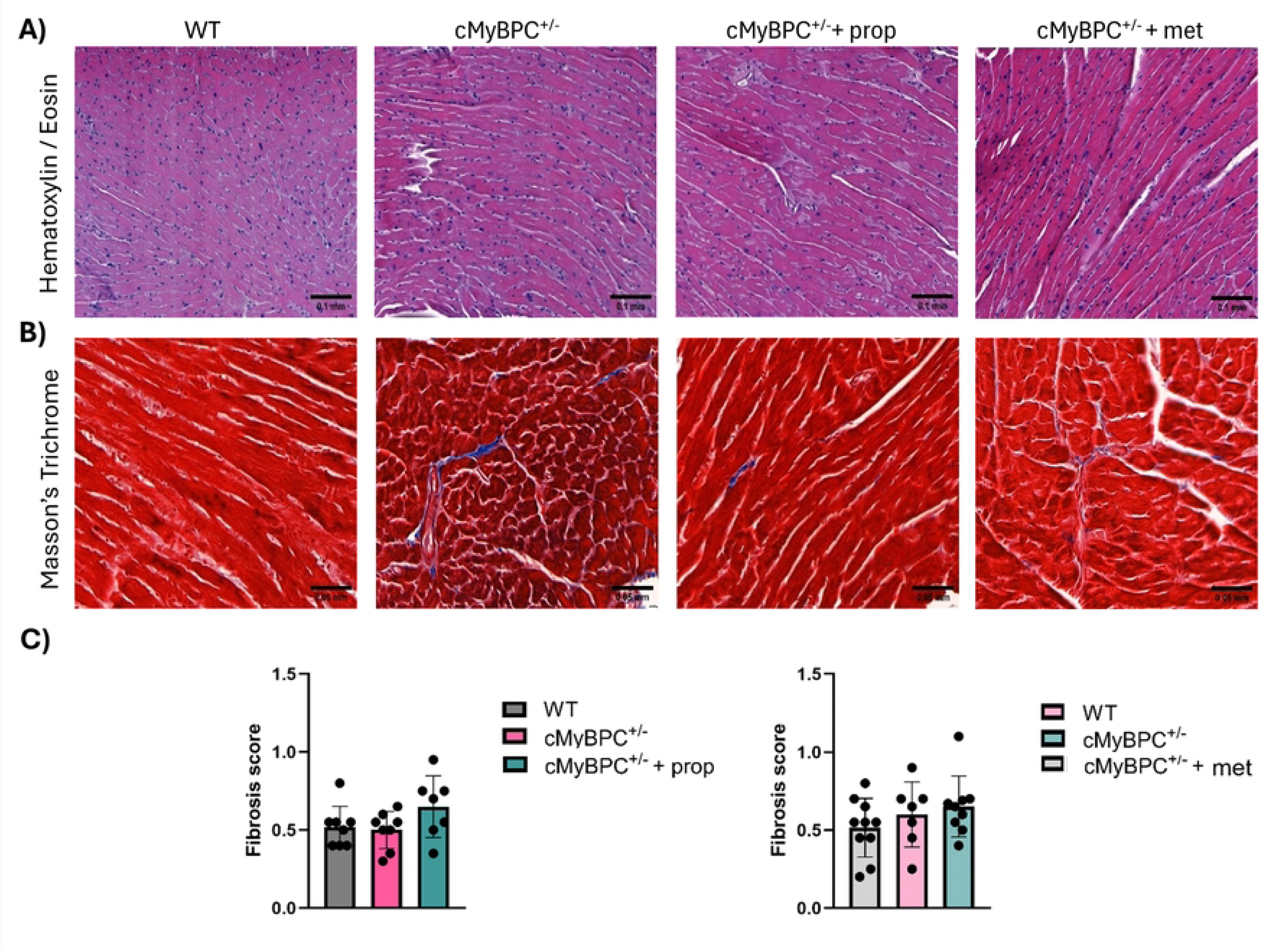
Representative images of cardiac sections from WT, wild type; cMyBPC^+/-^ + met, cMyBPC^+/-^ mice treated with metoprolol. cMyBPC^+/-^ + prop, cMyBPC^+/-^ mice treated with propranolol, stained with hematoxylin and eosin (A) (scale bar = 0.1 mm) or Masson’s trichrome (B) (scale bar = 0.05 mm). (C) Graphs show interstitial fibrosis (0 = no fibrosis, 1 = slight, 2 = moderate, 3 = severe) presented as a mean value per individual.

Another histopathological hallmark of HCM is interstitial fibrosis, where collagen fibres expand and replace the cardiomyocytes. Using Masson’s trichrome staining, the abundance of fibrous tissue was scored using a semi-quantitative analysis. The staining showed a small amount of interstitial collagen in samples from cMyBPC^+/-^ mice in Prop-group, often extending from vessels (Fig 3, lower panel). While not significant, the amount of fibrosis seemed to be slightly higher in cMyBPC^+/-^+prop group. Similarly, in Met-group, fibrosis was somewhat more abundant in cMyBPC^+/-^+met mice. No large fibrotic areas were found in either of the cMyBPC^+/-^mice.

### Expression of HA synthases was elevated in cMyBPC^+/-^mice

Hyaluronan is synthesized at the cell membrane by hyaluronan synthases (HAS1, HAS2, HAS3). In Met-group, no changes in expression of HA synthases were detected. In Prop-group, the expression of *Has1* was unaltered in both treated and untreated cMyBPC^+/-^ mice. However, the expression of *Has2* was increased in cMyBPC^+/-^ mice, whereas in cMyBPC^+/-^ +prop mice, *Has2* expression remained at WT levels. For *Has3* this trend was reversed – cMyBPC^+/-^+prop mice show significant increase in *Has3* expression in comparison with WT mice. In cMyBPC^+/-^ mice, *Has3* levels were unchanged (Table 4).

**Table 4.** Gene expression by RT-qPCR.

|  | Met-group |  |  | Prop-group |  |  |
| --- | --- | --- | --- | --- | --- | --- |
|  | WT | cMyBPC<sup>+/-</sup> | cMyBPC<sup>+/-</sup> + met | WT | cMyBPC<sup>+/-</sup> | cMyBPC<sup>+/-</sup> + prop |
| **HAS1** | 1.12 ± 0.5 | 1.21 ± 0.7 | 1.33 ± 0.7 | 0.95 ± 0.2 | 1.62 ± 1.4 | 1.24 ± 0.7 |
| **HAS2** | 1.03 ± 0.3 | 1.05 ± 0.2 | 1.14 ± 0.2 | 1.01 ± 0.2 | 1.37 ± 0.2* | 0.96 ± 0.2<sup>++</sup> |
| **HAS3** | 1.04 ± 0.3 | 0.99 ± 0.4 | 0.91 ± 0.2 | 1.04 ± 0.2 | 1.16 ± 0.2 | 1.61 ± 0.3*** <sup>++</sup> |
| **HYAL1** | 1.01 ± 0.2 | 0.96 ± 0.2 | 1.01 ± 0.1 | 1.02 ± 0.2 | 0.88 ± 0.2 | 1.02 ± 0.1 |
| **HYAL2** | 1.01 ± 0.2 | 0.90 ± 0.2 | 1.05 ± 0.1 | 1.01 ± 0.1 | 0.98 ± 0.1 | 1.03 ± 0.1 |
| **TMEM2** | N/A | N/A | N/A | 0.99 ± 0.4 | 1.06 ± 0.1 | 1.31 ± 0.1 |
| **CEMIP** | N/A | N/A | N/A | 1.09 ± 0.5 | 1.51 ± 0.6 | 0.86 ± 0.3 |
| **GLUT4** | 1.01 ± 0.2 | 1.34 ± 0.3 | 0.96 ± 0.1<sup>+</sup> | 1.00 ± 0.1 | 0.93 ± 0.1 | 1.07 ± 0.1 |

Data displayed as mean ± SD. WT, wild type; cMyBPC<sup>+/-</sup>+ met, cMyBPC<sup>+/-</sup> mice treated with metoprolol; cMyBPC<sup>+/-</sup> + prop, cMyBPC<sup>+/-</sup> mice treated with propranolol. Significance measured by one-way ANOVA with Tukey's post-hoc test. Significance cMyBPC<sup>+/-</sup> vs WT: \*P<0.05, \*\*P<0.01, \*\*\*P<0.001. cMyBPC<sup>+/-</sup> vs cMyBPC<sup>+/-</sup>+prop/met: <sup>+</sup>P<0.05, <sup>++</sup>P<0.01. N/A, not measured due to lack of material.

### Cell-surface hyaluronidases were affected by propranolol treatment

Since we observed changes in expression of genes involved in HA synthesis in Prop-group, we continued to investigate genes connected to HA degradation. The expression of the hyaluronidases *Hyal1* and *Hyal2* was on the same level in WT and cMyBPC^+/-^ mice in both Met-group and Prop-group, regardless of treatment.

In Prop-group, the expression of *Cemip* was decreased in cMyBPC^+/-^+prop mice in comparison with cMyBPC^+/-^ mice (approximately 57% decrease, P=0.0549), reaching similar expression level as in WT mice. Contrastingly, the expression of *Tmem2* was increased in cMyBPC^+/-^+ prop mice compared to untreated cMyBPC^+/-^ mice (approximately 18% increase, P=0.073) (Table 4). Expression of these genes was not measured in Met-group due to lack of material.

### Β-blocker treatment influenced metabolic pathways in the heart

Next, we investigated other genes that could be affected by *Mybpc3* mutations or β-blocker treatment, namely genes connected to glycolysis, fatty acid oxidation and inflammation. However, not many striking differences were detected.

In cMyBPC^+/-^ mice in Met-group, we observed an increase in the glucose transporter 4 (*Glut4*, also known as *Slc4a2*) expression. In contrast, in cMyBPC^+/-^+met mice, the expression of *Glut4* was significantly lower compared to cMyBPC^+/-^ mice, approximately on the same level as WT (Table 4). However, no significant changes were detected in *Glut4* expression in Prop-group.

### Cardiomyocyte size correlated with hyaluronan abundance in hearts of cMyBPC^+/-^ mice

To assess the amount of HA in the samples, sections of cardiac tissue were stained using an HA probe and percentage of positive staining was quantified in 16+16 tiles in pre-defined high c[HA] areas and low c[HA] areas. Fig 4 shows representative images of HA staining in low and high c[HA] areas, as well as an example of tilling on a cMyBPC^+/-^ sample.

**Fig 4.**
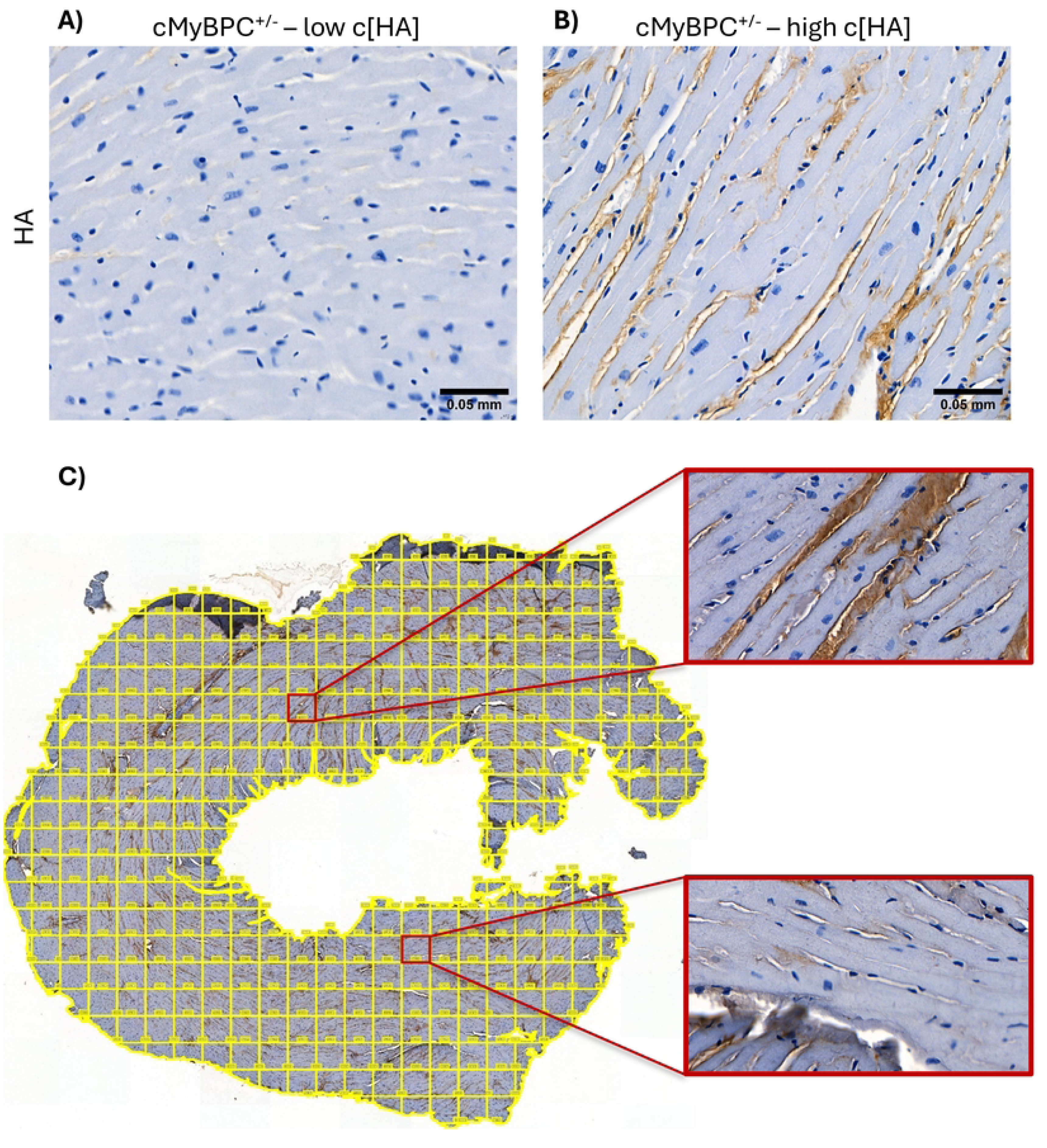
Representative images of HA staining showing a high c[HA] area (A) and a low c[HA] area (B) (scale bar = 0.05mm) on a left ventricle from a cMyBPC^+/-^ mouse. (C) Example of tiling for quantification of HA staining on a cMyBPC^+/-^ + prop heart, and an example of high c[HA] area (up), and a low c[HA] area (down). The cutoff for an area to be classified as low c[HA] was 5% of HA positive staining. For each type of area, the percentage of positive staining was quantified for 16 tiles and then averaged for the whole sample.

The results show that in high c[HA] areas, the amount of HA present is generally higher in cMyBPC^+/-^ mice compared to WT mice; in Prop-group, this difference is highly significant, whilst in the Met-group the data merely trend in this direction (Fig 5 left). The β-blocker treatment did not significantly affect HA abundance, regardless of the β-blocker used.

**Fig 5.**
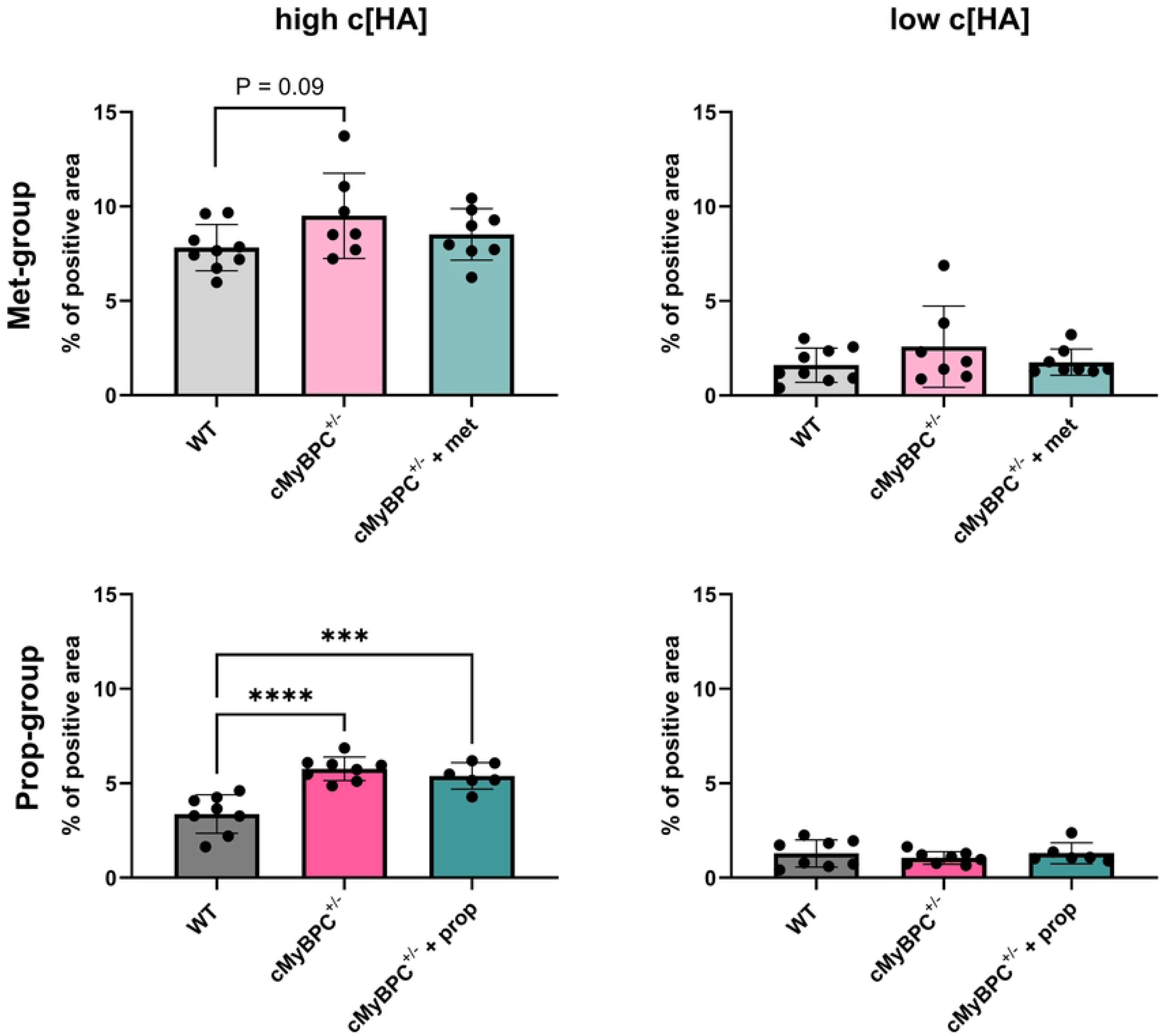
Average positive HA staining in either high c[HA] areas or low c[HA] areas. WT, wild type; cMyBPC^+/-^ + met, cMyBPC^+/-^ mice treated with metoprolol; cMyBPC^+/-^ + prop, cMyBPC^+/-^ mice treated with propranolol. n=7-10 mice per group, 16 tiles/sample. Data displayed as mean ± SD, significance was measured using one-way ANOVA with Tukey’s post-hoc test. ***P<0.001, ****P<0.0001.

In low c[HA] areas, the HA levels were comparable across all groups - less than 5% in all samples in both Met-group and Prop-group (Fig 5, right). This supports our hypothesis that hearts of cMyBPC^+/-^ mice contain patches of pathological tissue that produces overabundance of HA, as well as areas with healthy non-hypertrophic cardiomyocytes.

Lastly, we focused on cardiomyocyte size in the same pre-selected areas. The cell width was measured manually on longitudinally cut cells. The average cell width in high c[HA] areas was significantly larger in cMyBPC^+/-^ mice in comparison to WT mice in both Met-group and Prop-group (Fig 6, left). The β-blocker treatment did not seem to affect the cardiomyocyte hypertrophy, regardless of the β-blocker used. In low c[HA] areas, cell width was alike in all groups, between 10-13 µM, which corresponds with the levels of HA staining in these regions, which were alike in all groups as well (Fig 6, right). It is worth noting that for the control group, the cell size is nearly identical in both high c[HA] and low c[HA] areas. The data shows that cMyBPC^+/-^ mice, regardless of β-blocker treatment, have hypertrophied cardiomyocytes exclusively in areas rich in HA.

**Fig 6.**
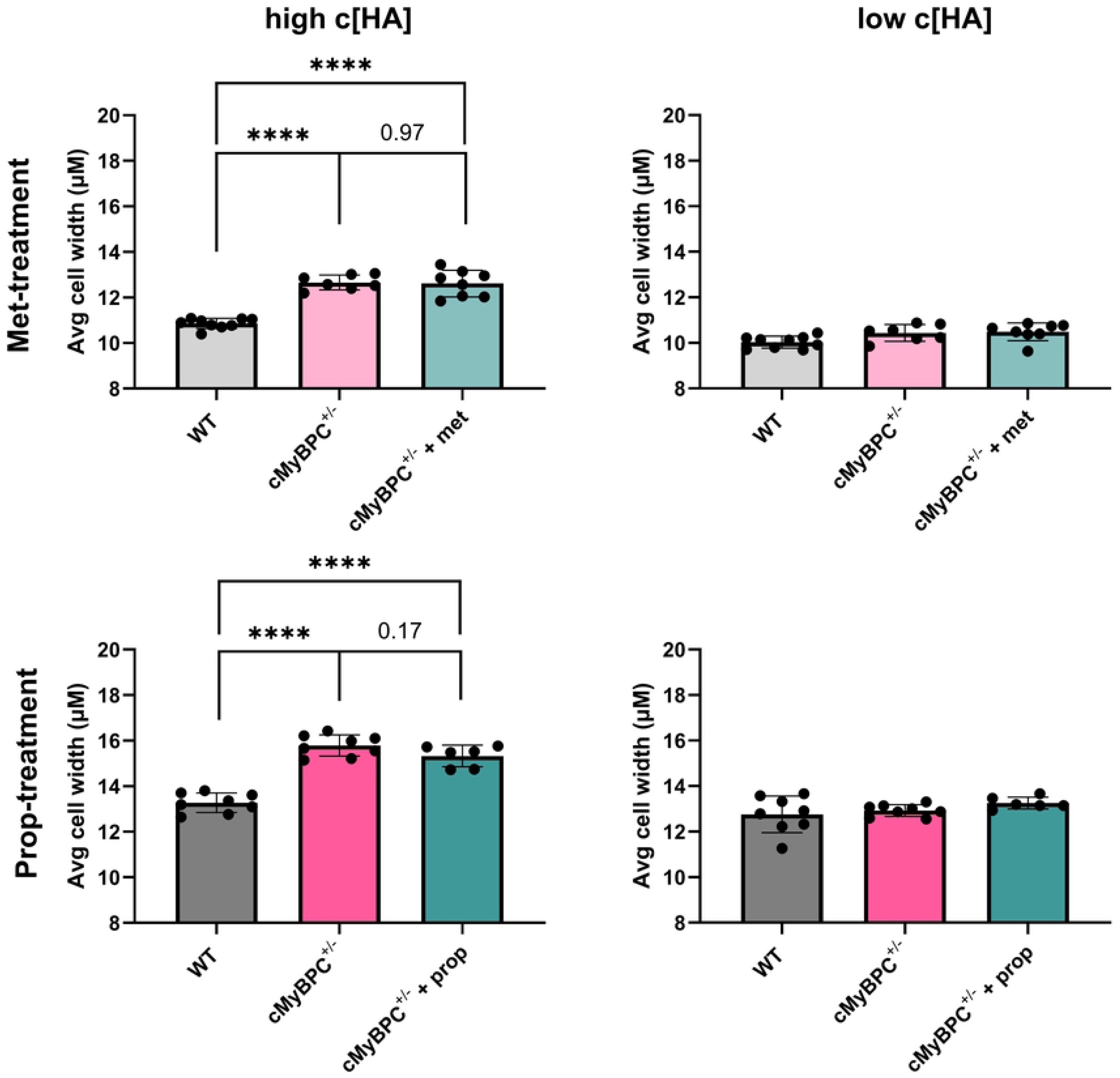
Average cardiomyocyte width in either high c[HA] areas or low c[HA] areas. WT, wild type; cMyBPC^+/-^ + met, cMyBPC^+/-^ mice treated with metoprolol; cMyBPC^+/-^ + prop, cMyBPC^+/-^ mice treated with propranolol. n=7-10 mice per group, n=50 cells/sample. Data displayed as mean ± SD, significance was measured using one-way ANOVA with Tukey’s post-hoc test. ***P<0.001, ****P<0.0001; ns, non-significant.

Furthermore, cell width and HA abundance were positively correlated (Pearson corr., r=0.489, P=0.02) in all mice, demonstrating a connection between these two features. The increase of cell width in cMyBPC^+/-^+ prop mice was also positively correlated with the expression of *Has3* (Pearson corr., r=0.776, P=0.04).

## Discussion

In this study, we investigated the connection between HA, glucose uptake and cardiomyocyte size in cMyBPC^+/-^ mice, as well as the effect of two β-blockers, metoprolol and propranolol, on HA production and hypertrophy when administered from early age (Fig 7). Metoprolol and propranolol are two of the most frequently used β-blockers. Both drugs are classed as non-vasodilating β-blockers, with different targets – propranolol is a non-selective β-1,2 antagonist, while metoprolol selectively affects β-1 receptors. Non-vasodilating β-blockers lower heart rate and contractility. Moreover, they have known detrimental effects on insulin sensitivity and glucose uptake and are associated with an increased risk of developing diabetes [23]. These effects on glucose usage originate from unopposed α-1-adrenergic activity as well as decreased blood flow to the peripheries [23,28].

**Fig 7.**
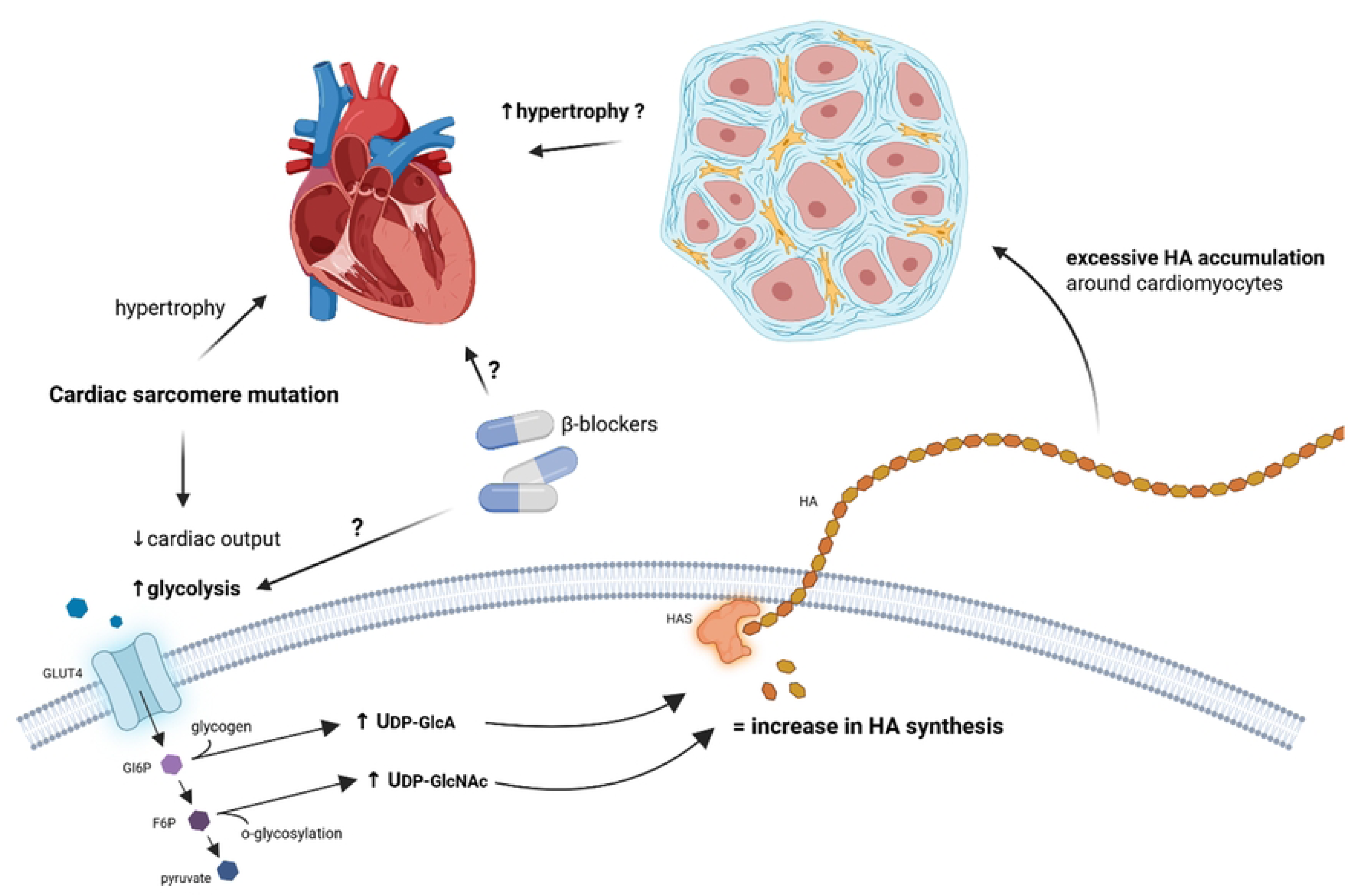
Summary of the main hypothesis and aims in the present study. In HCM, the heart experiences increased energy demands due to a mutation in cardiac sarcomere genes, such as cMyBPC haploinsufficiency. To fulfil these energy requirements, the heart starts primarily using glycolysis to make energy. Glycolytic intermediates can be used in side-pathways that give rise to HA precursors, UDP-GlcA and UDP-GlcNAc, resulting in increased HA production. Excessive HA accumulation in the extracellular space can disrupt correct cardiac function and contribute to hypertrophy. In this study, we investigated the connection between HA metabolism, glycolysis and cardiomyocyte hypertrophy using β-blockers known to disrupt glucose uptake.

We demonstrated that hypertrophied cardiomyocytes are solely located in areas with high HA abundance, while in areas with low amount of HA, the cardiomyocyte size remains normal. Indeed, cell width and HA abundance are positively correlated. In cMyBPC^+/-^+ prop mice, we found a connection between increase of HA concentration and the expression of *Has3*, showing that propranolol does affect HA synthesis, presumably via increasing *Has3* expression, while normalizing the expression of *Has2*. However, cardiomyocyte size itself was unaffected by the β-blocker treatment. Although cMyBPC^+/-^ mice were used for our research to better mimic human disease, complete knockout of the *Mybpc3* gene might have been able to show a stronger hypertrophic phenotype and subsequently a stronger effect of the treatment [29].

In the present study, we developed a technique for quantifying the amount of HA in cardiac sections. Dividing each sample into tiles helped attenuate individual differences between samples. Furthermore, we identified areas with high and low c[HA] levels and measured the percentage of positive HA staining in each. Quantifying HA in random tiles around the whole tissue section would diminish the difference between the groups, as areas with very low staining and very high staining would mix. We found that in high c[HA] areas, the amount of HA is significantly elevated in both untreated and treated cMyBPC^+/-^ mice in comparison to WT. In low c[HA] areas, the amount of HA remained at similar level in all groups. These results support the hypothesis that HCM is a ’patchy’ disease – meaning that the HCM heart comprises of mostly healthy normally-sized cardiomyocytes, which are interspaced with patches of abnormal hypertrophied cells [15].

RT-qPCR assays were performed to identify genes in the HA metabolism that were either affected by the *Mybpc3* mutation or by β-blocker treatment. RNA was extracted from whole tissue samples, meaning that hypertrophied cardiomyocytes were mixed with healthy cardiomyocytes and analysed in bulk. It is plausible that altered pathological expression of some genes is diluted by normal gene expression from healthy cells; meaning that transcriptional changes in the hypertrophic cardiomyocytes were masked by the more abundant healthy cardiomyocytes, and only severe changes in expression were detectable. Spatial transcriptomics would therefore be helpful in future experiments, to be able to distinguish between healthy and pathological tissue within the same heart.

In Prop-group, the expression of *Has2* was significantly upregulated in cMyBPC^+/-^ mice, suggesting increased HA production. In contrast, cMyBPC^+/-^+ prop mice had a significantly increased expression of *Has3*, and expression of *Has2* was maintained at WT level. In HCM, hearts rely primarily on glycolysis to produce energy; it is less oxygen demanding than creating ATP via β-oxidation, like healthy hearts would [13]. The increased amount of glucose is then utilized in glycolysis, creating high numbers of glycolytic intermediates like glucose-6-phosphate or fructose6-phosphate [30]. Instead of continuing via glycolysis into the TCA cycle, these intermediates can turn into HA precursors, UDP-glucuronic acid (UDP-GlcA) and UDP-N-acetyl-glucosamine (UDP-GlcNAc). Specifically, glucose-6-phosphate is converted into glucose-1-phosphate and UDP-glucose. UDP-glucose dehydrogenase then turns UDP-glucose into UDP-GlcA. On the other hand, UDP-GlcNAc is synthesized via the hexosamine pathway, starting with fructose-6-phosphate. Hexosamine pathway combines glutamine, acetyl-CoA and uridine-5’-triphosphate, resulting in the end product, UDP-GlcNAc [31,32]

HA synthesis depends on the availability of these UDP-precursors, and, by extension, on the availability of glucose. Up to 5% of all glucose molecules entering glycolysis are shuttled into synthesis of HA precursors [21]. Interestingly, when glucose uptake was measured in cMyBPC^+/-^ +prop mice with FDG-PET, the data formed two distinct clusters; one with high uptake, one with low uptake, with no data at the mean. This result does not correlate with the data distribution in results of HA positive staining, cell size nor gene expression. It is possible that glucose uptake was influenced by propranolol treatment, however, sensitivity to the β-blocker’s effect on glucose could vary in each individual.

When there is little glucose present in the cell, the concentration of UDP-precursors will be very low as well, resulting in changes in HA production. The activity of the individual HAS isoforms depends primarily on the cellular pool of UDP-GlcNAc and UDP-GlcA; while HAS2’s and HAS1’s activity increases with concentration of UDP-sugars, HAS3 is capable of HA synthesis at high speed even with minimal substrates available [33].

The elevated expression of *Has3* in cMyBPC^+/−^ mice treated with propranolol strongly suggests that HAS3 may serve as a compensatory mechanism for the reduced activity of HAS2. This indicates a critical functional redundancy within HA synthesis pathways. Regardless of how *Has2* expression or HAS2 enzymatic activity is decreased, HAS3 could assume a pivotal role in upholding HA production essential for cardiac function. The importance of this mechanism is underscored by previous findings showing that *Has2* knockout models are non-viable; absence of HAS2 leads to severe cardiac malformations and embryonic lethality [37]. Consequently, HAS3 may represent an evolutionary safeguard designed to ensure sufficient HA synthesis, even under conditions where HAS2 activity is compromised. However, HAS3 tends to produce slightly shorter HA molecules than HAS2 [34]; it is not known how this change influences the heart. Altogether, these results indicate that both the *Mybpc3* mutation and propranolol treatment affect HA metabolism, possibly by influencing glycolysis or glucose levels.

Surprisingly, in Met-group, no changes in expression of *Has* genes were observed (Table 4). The discrepancies between propranolol and metoprolol treated animals were presumed, due to difference in dosage as well as receptors affected. Propranolol, that affects both β1 and β2 receptor, was administered in a high dose of 37mg/kg/day, while metoprolol, a specific β1 inhibitor, had a dosage of 30mg/kg/day. Unexpectedly, no changes in expression of HA metabolism genes were detected in the cMyBPC^+/-^ mice in Met-group, whereas in cMyBPC^+/-^mice in Prop-group, the expression of *Has2* was elevated. This may be due to individual differences between the mice in Met-group and Prop-group, due to heterogeneity – which is a key aspect of this disease, or due to a compensatory effect from the remaining intact *Mybpc3* gene allele. Since the treatment was administered via drinking water over the course of approximately 8 months, it is unlikely that these differences are due to drinking habits of the mice. Interestingly, it has been reported that rats given propranolol in drinking water have a lower water intake than control rats [35].

In a previous study from our lab, aorta-ligated rats, which develop pressure-originated cardiac hypertrophy, showed a six-fold increase in the expression of *Has1* and *Has2* at day 1 post operation. Over the course of 42 days, *Has2* dropped to basal levels, while *Has1* dropped from six- to three-fold increase above normal level [16]. On the contrary, in human myectomy samples, we detected a decrease in expression of *HYAL2* and *HAS2*, and upregulation of *HAS3*; a similar pattern to what we observed in this study in cMyBPC^+/-^+ prop mice. This similarity between gene expression in mice in this study and human patients from the previous study could be explained by the fact that several patients were being treated with β-blockers [36]. To our knowledge, the exact mechanism in which β-blockers affect cardiac muscle cells and HA synthesis is unknown; however, in asthmatic airway smooth muscle cells, β2 antagonists have been observed to reverse pathological changes in HA metabolism when used together with steroids [37].

Depolymerisation of hyaluronan begins at the cell membrane, where the large molecules are cleaved into shorter fragments and then internalized. Recently, new cell surface hyaluronidases have been described – TMEM2 and CEMIP. While their function in mice has not yet been completely elucidated, they both can degrade HA into shorter fragments, which can then contribute to inflammation [38–41]. Surprisingly, human TMEM2 lacks hyaluronidase function, and serves instead as a regulator of CEMIP and HAS2 [38]. Even though we saw no changes in expression of traditional hyaluronidases, *Hyal2* and *Hyal1*, both *Cemip* and *Tmem2* were influenced by propranolol treatment. In cMyBPC^+/-^+ prop mice, the expression of *Cemip* was around 57% lower than in cMyBPC^+/-^ mice (P=0.0549). Similarly, expression of *Tmem2* was slightly elevated in cMyBPC^+/-^+ prop mice – approximately 18% higher than in cMyBPC^+/-^mice (P=0.073). This could indicate that HA degradation pathway is affected by propranolol. In our previous research, we have shown that hearts of both HCM patients and rats with pressure-derived cardiac hypertrophy contain high amounts of fragmented LMW HA [16,20]. It is possible that the concentration of LMW HA fragments increases in pathological conditions due to changes in activity or expression of HA degrading molecules, such as CEMIP or TMEM2.

We have also investigated the expression levels of genes related to glycolysis. We found that not the *Mybpc3* mutation itself, but treatment with metoprolol affects glucose uptake. The expression of *Glut4* was significantly downregulated in cMyBPC^+/-^+ met mice. Despite being applied in a higher dose, propranolol showed no effect on *Glut4* expression (Table 4). Latest research suggests that β2 receptor activity, which is blocked by propranolol, is required for GLUT4-mediated glucose uptake [42]. The decreased expression of *Glut4* observed after metoprolol treatment might be the cause for previously reported increased plasma glucose after metoprolol usage in diabetic patients [43]. However, we did not observe any changes in blood glucose levels in cMyBPC^+/-^ mice, regardless of treatment.

Echocardiography was used to monitor cardiac function. The cMyBPC^+/-^ mouse has been previously described by Carrier *et al.* as exhibiting increased diastolic septum and posterior wall thickness, as well as increased left ventricle inner diameter in both systole and diastole, at 10-11 month of age [25]. In the present study, no similar changes were found. We could see an overall reduced cardiac output in cMyBPC^+/-^ mice without treatment in the first study. The reduced cardiac output was improved but not restored when treated with metoprolol. However, these results were not reproduced in the second study where mice were given propranolol.

Nevertheless, we report significant cardiomyocyte hypertrophy localized in a focal patch-like pattern in the hearts of cMyBPC^+/-^ mice. The key mechanism by which cMyBPC variants cause disease is probably haploinsufficiency [44,45]. The healthy cMyBPC allele is not able to generate sufficient cMyBPC protein content to carry out its normal function in the sarcomere. This discrepancy between our study and Carrier *et. al.* could therefore be explained by different amounts of the cMyBPC protein in the heart leading to diverse outcomes; additionally, HCM is known to be a highly heterogenous disease [47]. Furthermore, in the study by Carrier *et al.* [25], the anaesthetic regime as well as the scan-frequency used for echocardiography were different. The study by Carrier *et al.* does not describe how the cardiac wall thickness was measured [25]. In the present study, wall thickness was measured in M-mode with the papillary muscle as anatomical hallmark. It is possible that hypertrophy would have been detected if more areas were chosen for measurement. Therefore, caution should be taken when comparing studies and models used.

Moreover, we observed differences between mice in Met-group and Prop-group. These mice were developed and experimented on independently within the span of three years and could therefore display inter-generational differences. The breeding was continuing from a few couples of founders that were either wild type or the cMyBPC^+/-^ genotype. This, together with the fact that the model is heterozygous, can result in a broader range of variations then a homozygotic knockout would give.

The present study describes a novel connection between HA and cardiomyocyte hypertrophy in cMyBPC^+/-^ mice. We observed that HA synthesis is affected by *MyBPC* mutation as well as β-blocker treatment. Cardiomyocyte size is significantly increased in areas with high HA concentration. This could suggest that changes in glucose metabolism contribute to hypertrophy by modulating extracellular matrix, including hyaluronan. Interestingly, we observed several differences between mice in Met-group and Prop-group. In Prop-group, we demonstrated an increase in *Has2* expression that led to higher abundance of HA and enlargement of cells in areas with high c[HA]. On the contrary, there was no change in *Has2* expression in Met-group. However, we still observed an increase in cell width, suggesting that there might be several pathways affecting HA production. Propranolol’s effect on HAS3 and HAS2 could influence HA synthesis directly via these HA synthases. There could also be another, more indirect, pathway, where HA production is affected via an increase of HAS2 activity, but not expression, due to a large spillover of glycolytic intermediates into HA synthesis. Further studies will be needed to explore the intricacies of these pathways, as well as the exact role these changes play in the development of hypertrophy, e.g. using cultured cardiomyocytes or cardiac organoids.

## Acknowledgements

We acknowledge the Small Animal Research Imaging Facility (SARIF) at Umeå University for providing assistance in using the PET/CT and ultrasound systems.

We acknowledge Kristina Eriksson for her laboratory assistance.

“Fig 7. Summary of the main hypothesis and aims in the present study.” was created in BioRender. Hellman, U. (https://BioRender.com/bnuzyrp) is licensed under CC BY 4.0.

## Author contributions

UH - conceptualization, formal analysis, investigation, supervision, administration, writing - review & editing, funding

ME – conceptualization, formal analysis, investigation, supervision, administration, writing - review & editing

RN – formal analysis, funding, investigation, supervision, administration, writing - review & editing

AS – formal analysis, investigation, funding, methodology, writing – original draft, visualisation

SM – conceptualization, supervision, funding, writing - review & editing

ID – formal analysis, investigation, writing - review & editing

All authors have read and agreed to the published version of the manuscript.

## Institutional Review Board Statement

The animal study was conducted in accordance with the Declaration of Helsinki and approved by the Animal Review Board at the Court of Appeal for Northern Norrland in Umeå (A62/14, approved on 2014-08-26).

## Declaration of Generative AI

The authors declare that no generative AI tools were used during the creation of this work.

## Abbreviations

Abbreviations used in this manuscript in order of appearance:

HCM: hypertrophic cardiomyopathy
cMyBPC: cardiac myosin binding protein C
HA: hyaluronan
WT: wild type
Met: metoprolol
Prop: propranolol
PET: positron emission tomography
FDG: fluorodeoxyglucose
ROI: region of interest
c[HA]: concentration of hyaluronan
HAS1: hyaluronan synthase 1
HAS2: hyaluronan synthase 2
HAS3: hyaluronan synthase 3
UDP-GlcA: UDP-glucuronic acid
UDP-GlcNAc: UDP-N-acetylglucosamine
TMEM2: transmembrane protein 2
CEMIP: cell migration inducing hyaluronidase 1
GLUT4: glucose transporter 4

## References

1. Teekakirikul P, Zhu W, Huang HC, Fung E. Hypertrophic Cardiomyopathy: An Overview of Genetics and Management. Biomolecules. 2019 Dec 16;9(12):878.

2. Ommen SR, Ho CY, Asif IM, Balaji S, Burke MA, Day SM, et al. 2024 AHA/ACC/AMSSM/HRS/PACES/SCMR Guideline for the Management of Hypertrophic Cardiomyopathy: A Report of the American Heart Association/American College of Cardiology Joint Committee on Clinical Practice Guidelines. Circulation. 2024 June 4;149(23):e1239–311.

3. Spudich JA, Nandwani N, Robert-Paganin J, Houdusse A, Ruppel KM. Reassessing the unifying hypothesis for hypercontractility caused by myosin mutations in hypertrophic cardiomyopathy. EMBO J. 2024 Oct;43(19):4139–55.

4. Maron MS, Rowin EJ, Maron BJ. The Paradigm of Sudden Death Prevention in Hypertrophic Cardiomyopathy. Am J Cardiol. 2024 Feb 1;212S:S64–76.

5. Kochi AN, Vettor G, Dessanai MA, Pizzamiglio F, Tondo C. Sudden Cardiac Death in Athletes: From the Basics to the Practical Work-Up. Medicina (Kaunas). 2021 Feb 14;57(2):168.

6. Kim KA, Jung MH. Current and emerging medical and surgical therapy in hypertrophic cardiomyopathy. J Cardiovasc Imaging. 2025 Sept 24;33(1):13.

7. Monda E, Bakalakos A, Rubino M, Verrillo F, Diana G, De Michele G, et al. Targeted Therapies in Pediatric and Adult Patients With Hypertrophic Heart Disease: From Molecular Pathophysiology to Personalized Medicine. Circulation: Heart Failure. 2023 Aug;16(8):e010687.

8. Östman-Smith I. Beta-Blockers in Pediatric Hypertrophic Cardiomyopathies. Rev Recent Clin Trials. 2014 June;9(2):82–5.

9. Hill MG, Sekhon MK, Reed KL, Anderson CF, Borjon ND, Tardiff JC, et al. Intrauterine Treatment of a Fetus with Familial Hypertrophic Cardiomyopathy Secondary to MYH7 Mutation. Pediatr Cardiol. 2015 Dec;36(8):1774–7.

10. Ashrafian H, Redwood C, Blair E, Watkins H. Hypertrophic cardiomyopathy:a paradigm for myocardial energy depletion. Trends Genet. 2003 May;19(5):263–8.

11. Ranjbarvaziri S, Kooiker KB, Ellenberger M, Fajardo G, Zhao M, Vander Roest AS, et al. Altered Cardiac Energetics and Mitochondrial Dysfunction in Hypertrophic Cardiomyopathy. Circulation. 2021 Nov 23;144(21):1714–31.

12. Sequeira V, Bertero E, Maack C. Energetic drain driving hypertrophic cardiomyopathy. FEBS Lett. 2019 July;593(13):1616–26.

13. Stanley WC, Recchia FA, Lopaschuk GD. Myocardial substrate metabolism in the normal and failing heart. Physiol Rev. 2005 July;85(3):1093–129.

14. Al-Biltagi M, El Razaky O, El Amrousy D. Cardiac changes in infants of diabetic mothers. World J Diabetes. 2021 Aug 15;12(8):1233–47.

15. Varnava AM, Elliott PM, Sharma S, McKenna WJ, Davies MJ. Hypertrophic cardiomyopathy: the interrelation of disarray, fibrosis, and small vessel disease. Heart. 2000 Nov;84(5):476–82.

16. Hellman U, Hellström M, Mörner S, Engström-Laurent A, Aberg AM, Oliviero P, et al. Parallel up-regulation of FGF-2 and hyaluronan during development of cardiac hypertrophy in rat. Cell Tissue Res. 2008 Apr;332(1):49–56.

17. Hellman U, Malm L, Ma LP, Larsson G, Mörner S, Fu M, et al. Growth factor PDGF-BB stimulates cultured cardiomyocytes to synthesize the extracellular matrix component hyaluronan. PLoS One. 2010 Dec 21;5(12):e14393.

18. Hellström M, Engström-Laurent A, Mörner S, Johansson B. Hyaluronan and collagen in human hypertrophic cardiomyopathy: a morphological analysis. Cardiol Res Pract. 2012;2012:545219.

19. Hellman U, Mörner S, Engström-Laurent A, Samuel JL, Waldenström A. Temporal correlation between transcriptional changes and increased synthesis of hyaluronan in experimental cardiac hypertrophy. Genomics. 2010 Aug;96(2):73–81.

20. Lorén CE, Dahl CP, Do L, Almaas VM, Geiran OR, Mörner S, et al. Low Molecular Mass Myocardial Hyaluronan in Human Hypertrophic Cardiomyopathy. Cells. 2019 Jan 29;8(2):97.

21. Caon I, Parnigoni A, Viola M, Karousou E, Passi A, Vigetti D. Cell Energy Metabolism and Hyaluronan Synthesis. J Histochem Cytochem. 2021 Jan;69(1):35–47.

22. Tavianatou AG, Caon I, Franchi M, Piperigkou Z, Galesso D, Karamanos NK. Hyaluronan: molecular size-dependent signaling and biological functions in inflammation and cancer. FEBS J. 2019 Aug;286(15):2883–908.

23. Fonseca VA. Effects of beta-blockers on glucose and lipid metabolism. Curr Med Res Opin. 2010 Mar;26(3):615–29.

24. Yang K, Li X, Qiu T, Zhou J, Gong X, Lan Y, et al. Effects of propranolol on glucose metabolism in hemangioma-derived endothelial cells. Biochem Pharmacol. 2023 Dec;218:115922.

25. Carrier L, Knöll R, Vignier N, Keller DI, Bausero P, Prudhon B, et al. Asymmetric septal hypertrophy in heterozygous cMyBP-C null mice. Cardiovasc Res. 2004 Aug 1;63(2):293–304.

26. Moak JP, Kaski JP. Hypertrophic cardiomyopathy in children. Heart. 2012 July;98(14):1044–54.

27. Muresan ID, Agoston-Coldea L. Phenotypes of hypertrophic cardiomyopathy: genetics, clinics, and modular imaging. Heart Fail Rev. 2021 Sept;26(5):1023–36.

28. Carnovale C, Gringeri M, Battini V, Mosini G, Invernizzi E, Mazhar F, et al. Beta-blocker-associated hypoglycaemia: New insights from a real-world pharmacovigilance study. British Journal of Clinical Pharmacology. 2021;87(8):3320–31.

29. van den Dolder FW, Dinani R, Warnaar VAJ, Vučković S, Passadouro AS, Nassar AA, et al. Experimental Models of Hypertrophic Cardiomyopathy: A Systematic Review. JACC: Basic to Translational Science. 2025 Apr 1;10(4):511–46.

30. Oikari S, Kettunen T, Tiainen S, Häyrinen J, Masarwah A, Sudah M, et al. UDP-sugar accumulation drives hyaluronan synthesis in breast cancer. Matrix Biology. 2018 Apr 1;67:63–74.

31. Kobayashi T, Chanmee T, Itano N. Hyaluronan: Metabolism and Function. Biomolecules. 2020 Nov 7;10(11):1525.

32. Zimmer BM, Barycki JJ, Simpson MA. Mechanisms of coordinating hyaluronan and glycosaminoglycan production by nucleotide sugars. Am J Physiol Cell Physiol. 2022 June 1;322(6):C1201–13.

33. Rilla K, Oikari S, Jokela TA, Hyttinen JMT, Kärnä R, Tammi RH, et al. Hyaluronan synthase 1 (HAS1) requires higher cellular UDP-GlcNAc concentration than HAS2 and HAS3. J Biol Chem. 2013 Feb 22;288(8):5973–83.

34. Itano N, Kimata K. Mammalian Hyaluronan Synthases. IUBMB Life. 2002;54(4):195–9.

35. The effect of propranolol on blood pressure and sympathetic nerve activity in the spontaneously hypertensive rat. Takeda, K., S. Sasaki, I. Kaimasu, M. Yoshimura, M. Nakagawa, H. Ijichi & R. D. Bunag; Jap. Heart J. 1980, 21, 559–559. [Internet]. [cited 2025 Sept 29]. Available from: https://www.jstage.jst.go.jp/article/ihj1960/21/4/21_4_563/_pdf/-char/en

36. Gennebäck N, Wikström G, Hellman U, Samuel JL, Waldenström A, Mörner S. Transcriptional Regulation of Cardiac Genes Balance Pro- and Anti-Hypertrophic Mechanisms in Hypertrophic Cardiomyopathy. Cardiogenetics. 2012 Dec;2(1):e5.

37. Papakonstantinou E, Klagas I, Karakiulakis G, Hostettler K, S’ng CT, Kotoula V, et al. Steroids and β2-agonists regulate hyaluronan metabolism in asthmatic airway smooth muscle cells. Am J Respir Cell Mol Biol. 2012 Dec;47(6):759–67.

38. Sato S, Miyazaki M, Fukuda S, Mizutani Y, Mizukami Y, Higashiyama S, et al. Human TMEM2 is not a catalytic hyaluronidase, but a regulator of hyaluronan metabolism via HYBID (KIAA1199/CEMIP) and HAS2 expression. J Biol Chem. 2023 June;299(6):104826.

39. Yamamoto H, Tobisawa Y, Inubushi T, Irie F, Ohyama C, Yamaguchi Y. A mammalian homolog of the zebrafish transmembrane protein 2 (TMEM2) is the long-sought-after cell-surface hyaluronidase. J Biol Chem. 2017 May 5;292(18):7304–13.

40. Liu Y, Hu G, Li Y, Kong X, Yang K, Li Z, et al. Research on the biological mechanism and potential application of CEMIP. Front Immunol. 2023 Aug 18;14:1222425.

41. Zha Y, Luo X, Ge Z, Zhang J, Li Y, Zhang S. KIAA 1199/CEMIP knockdown attenuates cardiac remodeling post myocardial infarction by activating TSP4 pathway in mice. Biochim Biophys Acta Mol Basis Dis. 2024 Dec;1870(8):167473.

42. Jovanovic A, Xu B, Zhu C, Ren D, Wang H, Krause-Hauch M, et al. Characterizing Adrenergic Regulation of Glucose Transporter 4-Mediated Glucose Uptake and Metabolism in the Heart. JACC: Basic to Translational Science. 2023 June;8(6):638–55.

43. Jacob S, Klimm HJ, Rett K, Helsberg K, Häring HU, Gödicke J. Effects of moxonidine vs. metoprolol on blood pressure and metabolic control in hypertensive subjects with type 2 diabetes. Exp Clin Endocrinol Diabetes. 2004 June;112(6):315–22.

44. van Dijk SJ, Dooijes D, dos Remedios C, Michels M, Lamers JMJ, Winegrad S, et al. Cardiac myosin-binding protein C mutations and hypertrophic cardiomyopathy: haploinsufficiency, deranged phosphorylation, and cardiomyocyte dysfunction. Circulation. 2009 Mar 24;119(11):1473–83.

45. Carrier L, Mearini G, Stathopoulou K, Cuello F. Cardiac myosin-binding protein C (MYBPC3) in cardiac pathophysiology. Gene. 2015 Dec 1;573(2):188–97.

46. Hellström M, Ericsson M, Johansson B, Faraz M, Anderson F, Henriksson R, et al. Cardiac hypertrophy and decreased high-density lipoprotein cholesterol in Lrig3-deficient mice. Am J Physiol Regul Integr Comp Physiol. 2016 June 1;310(11):R1045–1052.

47. Amundsen BH, Ericsson M, Seland JG, Pavlin T, Ellingsen Ø, Brekken C. A comparison of retrospectively self-gated magnetic resonance imaging and high-frequency echocardiography for characterization of left ventricular function in mice. Lab Anim. 2011 Jan;45(1):31–7.

48. Sadeghipour A, Babaheidarian P. Making Formalin-Fixed, Paraffin Embedded Blocks. In: Yong WH, editor. Biobanking: Methods and Protocols [Internet]. New York, NY: Springer; 2019 [cited 2025 Apr 29]. p. 253–68. Available from: 10.1007/978-1-4939-8935-5_22

49. Ericsson M, Steneberg P, Nyrén R, Edlund H. AMPK activator O304 improves metabolic and cardiac function, and exercise capacity in aged mice. Commun Biol. 2021 Nov 18;4(1):1306.

